# 2,4-Diaminothieno[3,2-*d*]pyrimidines, a new class of anthelmintic with activity against adult and egg stages of whipworm

**DOI:** 10.1101/254037

**Authors:** Frederick A. Partridge, Ruth Forman, Nicky J. Willis, Carole J.R. Bataille, Emma A. Murphy, Anwen E. Brown, Narinder Heyer-Chauhan, Bruno Marinič, Daniel J.C. Sowood, Graham M. Wynne, Kathryn J. Else, Angela J. Russell, David B. Sattelle

**Author notes:** Corresponding authors: (DBS), (KJE), (AJR). These authors contributed equally (Frederick A. Partridge, Ruth Forman, Nicky J. Willis, Carole J.R. Bataille).

## Abstract

The human whipworm *Trichuris trichiura* is a parasite that infects around 500 million people globally, with consequences including damage to physical growth and educational performance. Current drugs such as mebendazole have a notable lack of efficacy against whipworm, compared to other soil-transmitted helminths. Mass drug administration programs are therefore unlikely to achieve eradication and new treatments for trichuriasis are desperately needed. All current drug control strategies focus on post-infection eradication, targeting the parasite *in vivo*. Here we propose developing novel anthelmintics which target the egg stage of the parasite in the soil as an adjunct environmental strategy. As evidence in support of such an approach we describe the actions of a new class of anthelmintic compounds, the 2,4-diaminothieno[3,2-*d*]pyrimidines (DATPs). This compound class has found broad utility in medicinal chemistry, but has not previously been described as having anthelmintic activity. Importantly, these compounds show efficacy against not only the adult parasite, but also both the embryonated and unembryonated egg stages and thereby may enable a break in the parasite lifecycle.

**Author Summary:** The human whipworm, *Trichuris trichiura*, infects around 500 million people globally, impacting on their physical growth and educational performance. There are currently huge mass drug administration (MDA) programs aiming to control whipworm, along with the other major soil transmitted helminths, *Ascaris* and hookworm. However single doses of albendazole and mebendazole, which are used in MDA, have particularly poor effectiveness against whipworm, with cure rates less than 40%. This means that MDA may not be able to control and eliminate whipworm infection, and risks the spread of resistance to albendazole and mebendazole in the parasite population.

We are attempting to develop new treatments for parasitic worm infection, particularly focused on whipworm. We report the identification of a class of compounds, diaminothienopyrimidines (DATPs), which have not previously been described as anthelmintics. These compounds are effective against adult stages of whipworm, and also block the development of the model nematode *C. elegans*.

Our DATP compounds reduce the ability of treated eggs to successfully establish infection in a mouse model of human whipworm. These results support a potential environmental spray to control whipworm by targeting the infectious egg stage in environmental hotspots.

## Introduction

### Current anthelmintics

The benzimidazole anthelmintics albendazole and mebendazole are typically used to treat human whipworm infection but are compromised by lack of single-dose efficacy and the risk of resistance. Thus, existing drugs lack sufficient efficacy in mass drug administration (MDA) programs to adequately control or potentially eradicate whipworm. This is a major stumbling block in the WHO target to eliminate morbidity from soil transmitted helminthiases in children by 2020. The current approach for controlling soil-transmitted helminths such as *Trichuris* is mass drug administration of a single-dose of albendazole or mebendazole, typically repeated annually [1]. However for infection with *T. trichiura*, single doses of benzimidazoles lead to low cure rates, only 28% and 36% for albendazole and mebendazole respectively [2]. These cure rates are much lower than those of other major human soil-transmitted helminths, *Ascaris lumbricoides* and hookworm, demonstrating the need for improvements to therapy specifically targeting *Trichuris*. Indeed modelling studies have demonstrated that, due to these low cure rates, MDA with benzimidazoles does not interrupt whipworm transmission and thus cannot achieve eradication in many settings [3].

Furthermore, the experience from studies on veterinary parasites is that widespread usage of anthelmintics can lead to rapid development of resistance. The discovery of isolates of two species of gastrointestinal nematodes resistant to monepantel only four years after its introduction [4] underlies the real threat to control programmes imposed by emerging drug resistance. Indeed, the combination of MDA programs and low single-dose cure rates may facilitate the development of drug resistance in populations of human parasites. For example, resistance to benzimidazole drugs is caused by point mutations in β-tubulin. Such resistance mutations have been found in *T. trichiura* after mass drug administration [5], and have been found to increase in frequency after MDA. High frequency of resistance mutations in a population may be associated with lower egg-reduction rates after MDA [6]. Whilst there is no clear evidence yet of widespread anthelmintic resistance in human populations, identification of new drugs with novel mechanisms of actions is warranted to slow the development of drug resistance.

### *Trichuris* lifecycle

A *T. trichiura* infection becomes patent when adult female worms, embedded in the gut of the host, start to lay eggs. A single female worm can lay up to 20,000 eggs per day and these unembryonated eggs pass out with the faeces and embryonate in the soil. Development only proceeds further if the embryonated eggs are accidentally consumed via contact of the next host with contaminated food, water or soil. Once ingested, signals for hatching are received when the eggs reach the large intestine [7,8], the newly emerged first stage larvae invade the mucosal epithelium and development to the adult stage of the parasite occurs through a succession of larval moults. Importantly, even when active infections are successfully treated, hosts are constantly re-infected due to high levels of infective eggs present within the water and soil, which can remain viable for years.

Current anthelmintic programmes, including those targeting *Trichuris*, focus on post-infection eradication of existing infections. However, lifecycle stages outside of the host are also potential viable targets for small molecule drugs. Thus, both preventing egg embryonation and reducing the infectivity of embryonated eggs prior to ingestion offer targets that would break the parasite lifecycle.

### Screening *ex vivo T. muris* adults for new anthelmintic chemotypes

The mouse whipworm, *T. muris*, is a convenient model of the human whipworm as it can be grown routinely in the laboratory via infection of severe combined immune deficiency (SCID) mice. Screening *ex vivo* adult *T. muris* has been used to test the anthelmintic activity of a variety of compounds, including approved drugs with the potential for repurposing, and also plant extracts [9–11]. We recently reported a small molecule screen utilising an automated assay for assessment of the motility of *ex vivo T. muris* adults. This screen led to the identification of a class of molecules termed dihydrobenzoxazepinone (DHB) which demonstrated encouraging activity in this assay, as well as the ability to reduce *in vivo* infectivity of treated eggs [12]. Most of the active molecules identified from that screen belonged to the dihydrobenz[*e*][1,4]oxazepin-2(3*H*)-one chemotype, but interestingly one additional active was from a completely different structural class. Here we report the identification, synthesis and characterisation of a series of compounds belonging to this second chemotype, which has not previously been described as having anthelmintic activity, the 2,4-diamino thieno[3,2-*d*]pyrimidines (henceforth called diaminothienopyrimidines or DATPs).

## Materials and methods

### Ethics statement

All animal experiments were approved by the University of Manchester Animal Welfare and Ethical Review Board and performed under the regulation of the Home Office Scientific Procedures Act (1986) and the Home Office project licence 70/8127.

### *In vivo* culture of *Trichuris muris*

*T. muris* worms were cultured using severe combined immune deficiency (SCID) mice, at the Biological Services Facility at the University of Manchester. Male and female mice were infected with 200 infective embryonated *T. muris* eggs via oral gavage. Thirty-five days later, the mice were sacrificed. Adult *T. muris* were obtained from the intestine as previously described [12]. Worms were maintained in Roswell Park Memorial Institute (RPMI) 1640 media supplemented with penicillin (500 U/mL) and streptomycin (500 µg/mL) at approximately 37 °C and studied on the same day.

### *Ex vivo T. muris* adult maintenance for motility screen

Individual adult worms were added to wells containing 75 µL of RPMI-1640 medium, penicillin (500 U/mL), streptomycin (500 µg/mL) plus 1% v/v final concentration of dimethylsulfoxide (DMSO) or compound dissolved in DMSO. Plates were incubated at 37 °C, 5% CO_2_. Motility was determined after 24 hours.

### Automated motility assay

An automated system was used to quantify worm movement. An earlier version of this system has been previously described [13,14]. Two hundred frame movies of the whole plate were recorded at 10 frames per second and then motility determined by an algorithm based on thresholding pixel variance over time [15]. For the hit confirmation and expansion assays, library material was used at a final concentration of 100µM. Dose-response curves were calculated with the four factor log-logistic model using the R package *drc* [16] or using GraphPad Prism.

### Chemical synthesis

Thin layer chromatography (TLC) was performed on aluminium sheets coated with 60 F_254_ silica. All solvents are used anhydrous unless stated otherwise. NMR spectra were recorded on Bruker AV400 (400 MHz), Bruker AVII 500 (500 MHz) or AVIIIHD 600 (600 MHz) instruments in the deuterated solvent stated. All chemical shifts (δ) are quoted in ppm and coupling constants (*J*), which are not averaged, in Hz. Residual signals from the solvents were used as an internal reference using the stated deuterated solvent. Infrared spectra were recorded on a Perkin-Elmer 1750 IR Fourier Transform spectrophotometer using thin films on a diamond ATR surface (thin film). Only the characteristic peaks are quoted. Melting points were determined using a Stanford Research Systems EZ-Melt. Low resolution mass spectra (*m/z*) were recorded on an Agilent 6120 spectrometer and high resolution mass spectra (HRMS *m/z*) on a Bruker microTOF mass analyzer using electrospray ionization (ESI). Compounds were synthesised from commercially available starting materials, and fully characterised by Infrared (IR) Spectroscopy, Mass Spectrometry (ESI-MS, HRMS-ESI) and Nuclear Magnetic Resonance (^1^H and ^13^C NMR). Spectra supporting the synthesis of these compounds are provided in the S1 File.

#### *2-Chloro-N-(2-(chlorophenoxy)ethyl)thieno[3,2-d]pyrimidin-4-amine* (2a)

To a 20 mL microwave vial containing 2,4-dichlorothieno[3,2-*d*]pyrimidine (1.50 g, 7.32 mmol, 1.0 equiv,) in 1,4-dioxane (15 mL) at RT was added 2-(2-chlorophenoxy)ethylamine (1.26 g, 7.32 mmol, 1.0 equiv.) and *N*,*N*-diisopropylethylamine (2.5 mL, 14.64 mmol, 2.0 equiv.) under an argon atmosphere. The vessel was sealed and the reaction heated at 80 °C for 3 hours. The mixture was cooled to RT, concentrated *in vacuo* and the crude residue was purified by flash column chromatography (silica gel) to afford the title compound as an off-white solid (1.59 g, 64%).

mp = 118–119 °C; R*_f_* = 0.2; ν_max_ (film)/cm^-1^ = 3398m (NH), 3088w (CH), 2970w (CH), 1586s (arom.), 1539m (arom.), 1483m (arom.); ^1^H NMR (500 MHz, CDCl_3_) δ 7.75 (1H, d, *J* = 5.4 Hz), 7.39 (1H, dd, *J* = 8.0, 1.6 Hz), 7.36 (1H, d, *J* = 5.4 Hz), 7.23 (1H, ddd, *J* = 8.3, 7.5, 1.6 Hz), 7.02 (1H, dd, *J* = 8.4, 1.4 Hz), 6.96 (1H, apparent td, *J* = 7.9, 1.3 Hz), 5.79 (1H, t, *J* = 5.4 Hz), 4.30 (2H, t, *J* = 5.0 Hz) 4.13 (2H, apparent q, *J* = 5.4 Hz); ^13^C NMR (500 MHz, CDCl_3_) δ 161.4, 158.1, 157.6, 153.9, 132.6, 130.5, 128.1, 124.9, 123.5, 122.6, 114.7, 114.3, 68.1, 40.5; LRMS (ESI^-^) calculated for [C_14_H_11_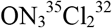S-H]^-^ = 338.0, found 337.9, [M-H]^-^, 100%, calculated for [C_14_H_11_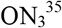Cl^37^Cl^32^SH]^-^ = 340.0, found, 339.9 [M-H]^-^, 60%; HRMS (ESI^+^) calculated for [C_14_H_11_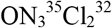S+H]^+^ = 340.0073, found 340.0071, [M+H]^+^.

#### *2-Chloro-N-(2-phenoxyethyl)thieno[3,2-d]pyrimidin-4-amine* (2b)

To a 20 mL microwave vial containing 2,4-dichlorothieno[3,2-*d*]pyrimidine (1.0 g, 5.0 mmol, 1.0 equiv.) in 1,4-dioxane (10 mL) at RT was added 2-phenoxyethylamine (0.6 mL, 5.0 mmol, 1.0 eq.) and *N*,*N*-diisopropylethylamine (1.7 mL, 10.0 mmol, 2.0 equiv.) under an argon atmosphere. The vessel was sealed and the reaction heated at 80 °C for 3 hours. The mixture was cooled to RT, concentrated *in vacuo* and the crude residue was purified by flash column chromatography (silica gel) to afford the title compound as an off-white solid (1.23 g, 80%).

mp = 115.5–116.9 °C; R*_f_* = 0.5 (EtOAc: Petroleum; 1:4); ν_max_ (film)/cm^-1^ = 3228w (NH), 3041w (CH), 2962w (CH), 1597s (arom.), 1581s (arom.), 1533m (arom.), 1511m (arom.), (arom.), 1496m (arom.), 1469m (arom.), 1434m (arom.); ^1^H NMR (500 MHz, CDCl_3_) δ 7.72 (1H, d, *J* = 5.4 Hz), 7.34 (1H, d, *J* = 5.4 Hz), 7.32-7.28 (2H, m), 6.97 (1H, app t, *J* = 7.3 Hz), 6.96-6.92 (2H, m), 5.78 (1H, t, *J* = 4.8 Hz), 4.22 (2H, t, *J* = 5.1 Hz), 4.11-4.06 (2H, m); ^13^C NMR (126 MHz, CDCl_3_) δ 161.4, 158.4, 158.1, 157.5, 132.6, 129.7, 124.8, 121.5, 114.6, 114.1, 66.2, 40.9; LRMS (ESI^+^) calculated for [C_14_H_12_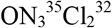S+H]^+^ = 306.0, found 306.0, [M+H]^+^, 100%, calculated for [C_14_H_12_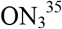Cl^37^Cl^32^S+H]^+^ = 308.0, found 308.0, [M+H]^+^, 40%, calculated for [C_14_H_12_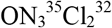S+Na]^+^ = 328.0, found 328.0, [M+H]^+^, 60%, calculated for [C_14_H_12_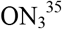Cl^37^Cl^32^S+Na]^+^ = 330.0, found 330.0, [M+H]^+^, 20%; HRMS (ESI^+^) calculated for [C_14_H_12_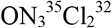S+H]^+^ = 306.0462, found 306.0462, [M+H]^+^.

### General Synthetic Procedure

To a 10 mL microwave vial containing 2-chlorothieno[3,2-*d*]pyrimidine (1.0 equiv) in ^*i*^PrOH (10 µL/mg chloride) at room temperature was added the requisite amine (10.0 equiv.) under an argon atmosphere. The vessel was sealed and the mixture heated at 100°C for 16-24 hours. The reaction was cooled to ambient temperature (RT), concentrated *in vacuo* and the crude residue was purified by flash column chromatography (silica gel).

#### *N2-Methyl-N4-(2-phenoxyethyl)thieno[3,2-d]pyrimidine-2,4-diamine* (3a, OX02925)

Following general procedure 1, the title compound was obtained from **2b** (600 mg, 1.96 mmol, 1.0 equiv.) and methylamine (2.0 M in THF, 9.8 mL, 19.6 mmol, 10.0 eq). Purification by flash column chromatography (MeOH:CH_2_Cl_2_; 1:49 v/v) followed by trituration with cold Et_2_O afforded the desired product as a pale yellow viscous oil (526 mg, 89%).

R*_f_* = 0.2 (MeOH:CH_2_Cl_2_; 1:49 v/v); ν_max_ (film)/cm^-1^ = 3418w (NH), 3232w (NH), 3038w (CH), 2936w (CH), 1585s (arom.), 1532s (arom.), 1508s (arom.), 1460s (arom.), 1405m (arom.); ^1^H NMR (500 MHz, CDCl_3_) δ 7.55 (1H, d, *J* = 5.4 Hz), 7.33-7.28 (2H, m), 7.15 (1H, d, *J* = 5.4 Hz), 7.00-6.96 (1H, m), 6.96-6.93 (2H, m), 5.16 (1H, brs), 4.83 (1H, brs), 4.21 (2H, t, *J* = 5.26 Hz), 4.02 (2H, m), 3.04 (3H, d, *J* = 5.04 Hz); ^13^C NMR (126 MHz, CDCl_3_) δ 161.9, 161.8, 158.5, 157.4, 130.4, 129.6, 124.1, 121.2, 114.5, 106.5, 66.6, 40.2, 28.7; LRMS (ESI^+^) calculated for [C_15_H_16_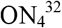S+H]^+^ = 301.1, found 301.1 [M+H]^+^, 100%; HRMS (ESI^+^) calculated for [C_15_H_16_ON_4_C^32^S+H]^+^ = 301.1119, found 301.1118 [M+H]^+^.

#### *N*4-(2-(2-Chlorophenoxy)ethyl)-N2-(2-methoxyethyl)thieno[3,2-d]pyrimidine-2,4-diamine (3b, OX02926)

Following general procedure 1, the title compound was obtained from **2a** (600 mg, 1.76 mmol, 1.0 equiv.) and 2-methoxyethylamine (1.5 mL, 17.6 mmol, 10.0 eq). Purification by flash column chromatography (MeOH:CH_2_Cl_2_; 1:49 v/v) followed by trituration with cold Et_2_O afforded the desired product as an off-white solid (380 mg, 57%).

mp = 69–97 °C (Et_2_O); R_*f*_ = 0.1 (MeOH:CH_2_Cl_2_; 1:49 v/v); ν_max_ (film)/cm^-1^ = 3424w (NH), 3304w (NH), 3076w (CH), 2949w (CH), 1606m (arom.), 1532s (arom.), 1476m (arom.), 1460m (arom.), 1444m (arom.), 1412m (arom.); ^1^H NMR (400 MHz, CDCl_3_) δ 7.56 (1H, d, *J* = 5.3 Hz), 7.39 (1H, dd, *J* = 7.9, 1.4 Hz), 7.23-7.21 (1H, m), 7.13 (1H, d, *J* = 5.3 Hz), 7.00-6.92 (2H, m), 5.29 (1H, s), 5.17 (1H, s) 4.27 (2H, t, *J* = 5.2 Hz), 4.05 (2H, apparent q, *J* = 5.4 Hz), 3.68-3.59 (4H, m), 3.40 (3H, s); ^13^C NMR (151 MHz, CDCl_3_) δ 161.8, 161.1, 157.3, 154.0, 130.5, 130.3, 127.8, 124.0, 123.3, 122.1, 114.3, 106.8, 71.6, 68.2, 58.7, 41.4, 39.9; LRMS (ESI^+^) calculated for [C_17_H_19_O_2_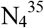Cl^32^S+H]^+^ = 379.1, found 379.1, [M+H]^+^, 100%, calculated for [C_17_H_19_O_2_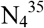Cl^32^S+Na]^+^ = 401.1, found 401.1, [M+Na]^+^, 10%; HRMS (ESI^+^) calculated for [C_17_H_19_O_2_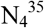Cl^32^S+H]^+^ = [M+H]^+^, 379.0990, found 379.0991 [M+H]^+^.

#### *N4-(2-(2-chlorophenoxy)ethyl)-N2-(2-methoxybenzyl)thieno[3,2-d]pyrimidine-2,4-diamine* (3c, OX03143)

Following general procedure 1, the title compound was obtained from **2a** (240 mg, 0.70 mmol, 1.0 eq.) and 2-methoxybenzylamine (0.92 mL, 7.0 mmol, 10.0 eq.). Purification by flash column chromatography (MeOH:CH_2_Cl_2_; 3:37 v/v) afforded the desired product (189 mg, 61%) as a thick pale yellow oil.

R_*f*_ = 0.4 (MeOH:CH_2_Cl_2_; 3:22 v/v); ν_max_ (film)/cm^-1^= 3424w (NH), 3247w (NH), 2935w (CH), 1587m (arom.), 1553 (arom.), 1487 (arom.), 1461 (arom.); ^1^H NMR (600 MHz, CDCl_3_) δ 7.54 (1H, d, *J* = 5.3 Hz), 7.38 (1H, dd, *J* = 8.1, 1.7 Hz), 7.36 (1H, s) 7.22 (1H, ddd, *J* = 9.5, 7.5, 1.6 Hz), 7.20 (1H, ddd, *J* = 8.2, 7.5, 1.6 Hz), 7.12 (1H, d, *J* = 5.3 Hz), 6.93 (1H, dt, *J* = 7.5, 1.5 Hz), 6.91 (1H, ddd, *J* = 8.1, 7.1, 1.1 Hz), 6.89 (2H, d, *J* = 7.9 Hz), 5.34 (1H, br), 5.29 (1H, t, *J* = 5.3 Hz), 4.68 (2H, d, *J* = 6.2 Hz) 4.20 (2H, t, *J* = 5.3 Hz), 4.04 (2H, apparent q, *J* = 5.3 Hz), 3.87 (3H, s); ^13^C NMR (151 MHz, CDCl_3_) δ 161.9, 161.2, 157.6, 157.4, 154.1, 130.5, 130.3, 129.1, 128.1 (x2), 127.8, 123.9, 123.3, 122.1, 120.3, 114.3, 110.1, 106.6, 68.2, 55.3, 41.4, 40.0; LRMS (ESI^+^) calculated for [C_22_H_21_O_2_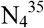Cl^32^S+H]^+^ = 441.1, found 441.2, [M+H]^+^, 100%; HRMS (ESI^+^) calculated for [C_22_H_21_O_2_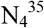Cl^32^S+H]^+^ = 441.1147, found 441.1142 [M+H]^+^.

#### *N4-(2-(2-chlorophenoxy)ethyl)-N2-methylthieno[3,2-d]pyrimidine-2,4-diamine* (3d, OX03147)

Following general procedure 1, the title compound was obtained from **2a** (237 mg, 0.70 mmol) and methylamine (2.0 M in THF) (3.5 mL, 7.0 mmol, 10 eq). Purification by flash column chromatography (MeOH:CH_2_Cl_2_; 1:19 v/v) afforded the desired product (218 mg, 93%) as a pale brown oil.

R_*f*_ = 0.4 (MeOH:CH_2_Cl_2_; 1:19 v/v); ν_max_ (film)/cm^-1^= 3247w (NH), 2940w (CH), 1588m (arom.), 1552m (arom.), 1510m (arom.), 1484m (arom.), 1461m (arom.), 1446m (arom.), 1406 (arom.); ^1^H NMR (600 MHz, CDCl_3_) δ 7.56 (1H, d, *J* = 5.3 Hz), 7.38 (1H, dd, *J* = 7.9, 1.7 Hz) 7.21 (1H, ddd, *J* = 8.2, 7.5, 1.7 Hz), 7.15 (1H, d, *J* = 5.3 Hz), 6.98 (1H, dd, *J* = 8.2, 1.5 Hz), 6.95 (1H, dd, *J* = 7.5, 1.5 Hz) 5.31 (1H, t, *J* = 5.5 Hz), 4.84 (1H, s) 4.27 (2H, t, *J* = 5.4 Hz), 4.06 (2H, apparent q, *J* = 5.5 Hz), 3.04 (3H, d, *J* = 5.0 Hz); ^13^C NMR (151 MHz, CDCl_3_) δ 162.0, 161.9, 157.5, 154.2, 130.7, 130.5, 128.0, 124.2, 123.6, 122.3, 114.6, 106.9, 68.5, 40.1, 28.9; LRMS (ESI^+^) calculated for [C_15_H_15_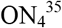S+H]^+^ = 335.1, found 335.0, [M+H]+, 100%, calculated for [C_15_H_15_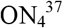S+H]^+^ = 337.1, found 337.0, [M+H]+, 30%, HRMS (ESI^+^) calculated for [C_15_H_15_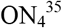S+H]^+^ = 335.0728, found 335.0725 [M+H]^+^, calculated for [C_15_H_15_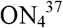S+H]^+^ = 337.0698, found 337.0695 [M+H]^+^.

### *C. elegans* growth assay

A mixed-stage *C. elegans* N2 population was obtained by liquid culture (20 °C) according to standard methods [17]. It was then bleached to obtain an egg population with 1.5 mL 4M NaOH, 2.4 mL NaOCl, 2.1 mL water, washed three times, and allowed to hatch in 50 mL S-basal buffer at 20 °C overnight to obtain a synchronised L1 population. For the growth assay, 49 µL of S-complete buffer and 1 µL of DMSO or DMSO plus compound were added to each well of 96-well plates. 50 µL of a worm suspension (approximately 20 synchronised L1 worms, 1% w/v *E. coli* HB101 in S-complete buffer) were then added to each well. Plates were incubated at 20 °C before imaging 5 days later. Worm movement was stimulated by inserting and removing a 96-well PCR plate into/from the wells of the assay plate, and then whole plate 200 frame movies were recorded at 30 frames per second. Growth was quantified as a correlate of movement using the same automated system described earlier, which estimates movement for each well by categorising pixels as imaging movement if their variance is greater the mean plus one standard deviation of the variances of all the pixels on the plate [15].

### Cytotoxicity testing

The mouse rectal epithelial cell line CMT-93 (LGC Promochem, Teddington, United Kingdom) was used for these studies. The WST-8 and neutral red cytotoxicity assays were performed as described [12]. Briefly, cells were cultured with test compounds, chlorpromazine positive control or DMSO alone (final compound concentrations of 0 to 100 µM) for 72 hours. The WST-8 assay was then carried out using the Cell Counting Kit – 8 (Sigma Aldrich # 96992) with an incubation time of 2 hours. This time was chosen according to the manufacturer’s instructions and was such that the absorbance of the WST-8 formazan dye was within the linear range of the microplate reader. Following this assay, the medium was exchanged, and the ability of the cells to take up the dye neutral red (concentration 33 µg/mL, incubation time 2 hours) was determined using a microplate reader (absorbance at 540 nm). Results were analysed using GraphPad Prism and fitted using a log-logistic model.

### *In vitro* and *in vivo* establishment of infection

100 infective embryonated eggs were incubated in deionised water with 1% v/v DMSO or test compounds at a final concentration of 100 µM in 1% v/v DMSO for 14 days at room temperature in the dark. Eggs were then washed and resuspended in deionised water. For *in vitro* hatching assays 100 eggs were added to 1 mL of *E. coli* bacterial culture grown in LB broth overnight at 37 °C shaking at 200 rpm. Egg-bacterial cultures were incubated for 24 hours at 37 °C, 5% CO_2_ and hatching determined following blinding by visual examination under a dissecting microscope. For *in vivo* hatching assays, 40 eggs were counted under a dissecting microscope and given to a SCID mouse in 200 µL water. At day 15 post-infection mice were culled and the number of L2 larvae present in the caecae and colon enumerated in a blinded manner under a dissecting microscope.

### Statistical analysis of *in vivo* establishment of infection data

The experiment was conducted in two ‘experimental batches’. For batch one there were 5 mice in each of the DMSO and **OX02926** groups. For batch two there were 9 mice in each of the DMSO and **OX02926** groups. The raw data (number of worms that established infection in each mouse) are shown separated by batch and treatment in the S2 Figure.

To analyse the data we used a two-way ANOVA (worm number ~ treatment * batch). This showed a significant effect of treatment [F(1,24) = 8.520, P = 0.00752]. It also showed a significant effect of batch [F(1,24) = 10.956, P = 0.00294]. There was no significant interaction between treatment and batch [F(1,24) = 0.296, P = 0.59153]. The significant effect of batch reflected that in both DMSO-and **OX02926**-treated groups, the number of worms that established infection was generally lower in mouse batch 1 than in batch 2 (S2 Figure). Variation in control worm establishment, which is commonplace in *Trichuris* infections due to natural variation in egg infectivity from a standardised egg number, was within expected ranges. We therefore took the approach of normalising each data point by dividing by the mean of the DMSO-treated group for that batch. This yielded the % batch normalised infection establishment.

We used a two-way ANOVA (% batch normalised infection establishment ~ treatment * batch) to analyse the data. There was a significant effect of treatment [F(1,24) = 9.569, P = 0.00497] but no effect of batch [F(1,24) = 0.083, P = 0.77618] or interaction [F(1,24) = 0.083 0.77618]. We therefore conducted a post-hoc Tukey HSD test which showed that infection establishment in the **OX02926-**treated group was significantly different from the DMSO-treated control group (P = 0.0050).

### Embryonation assay

One hundred unembryonated eggs were treated with water, 1% v/v DMSO in water or test compounds at a final concentration of 100 µM (unless stated) with 1% v/v DMSO, in the dark at 26 °C, either for 56 days or for shorter periods as described. Images were collected on an Olympus BX63 upright microscope using a 60x / 1.42 PlanApo N (Oil) objective and captured and white-balanced using an DP80 camera (Olympus) in monochrome mode through CellSens Dimension v1.16 (Olympus). Images were then processed and analysed using the image analysis platform Fiji [18].

### Data availability

Structures of resynthesized compounds have been deposited in the PubChem database with CID 49790760, 49790669, 46948320 and 49778268 and SID 348479445, 348479446, 348479447 and 348479448. Assay results for resynthesized compounds have been deposited in the PubChem database with assay ID 1259352 and 1259353.

## Results

### *Ex vivo T. muris* adult motility screen

We have recently described a small molecule screen for new anthelmintics, which used reduction or loss of motility of adult *ex vivo T. muris* as an endpoint for screening [12]. This screen was designed to identify compounds active on *Trichuris* as existing drugs are notably less efficacious against this nematode, and it is comparatively evolutionarily distant to nematodes typically screened in anthelmintic-discovery efforts, such as *H. contortus, M. incognita* and *C. elegans*. From this primary screen, we found 13 members of the dihydrobenzoxazepinone chemotype, which had not previously been shown to have anthelmintic activity.

In this report we describe the identification of a second new anthelmintic chemotype from this screen. A single 2,4-diaminothieno[3,2-*d*]pyrimidine (DATP) compound was found in the primary screen. This has been given the identifier **OX02926** (Fig 1a). We confirmed this activity in a secondary screen using the same source sample (DMSO solution containing 10 mM compound), and also tested a number of structurally-related compounds from our small molecule collection using the same assay (Fig 1b). The rationale for this was to gain greater confidence in the screening hit and also to explore the activity of “near-neighbour” molecules with the same core 2,4-diaminothieno[3,2-d]pyrimidine structure, which could support the early development of the series. The hit expansion process led to the identification of three further active molecules in this series **OX02925, OX03143** and **OX03147** (Fig 1c). Two structurally-related compounds were however not active in this assay (Fig 1d).

**Fig 1.**
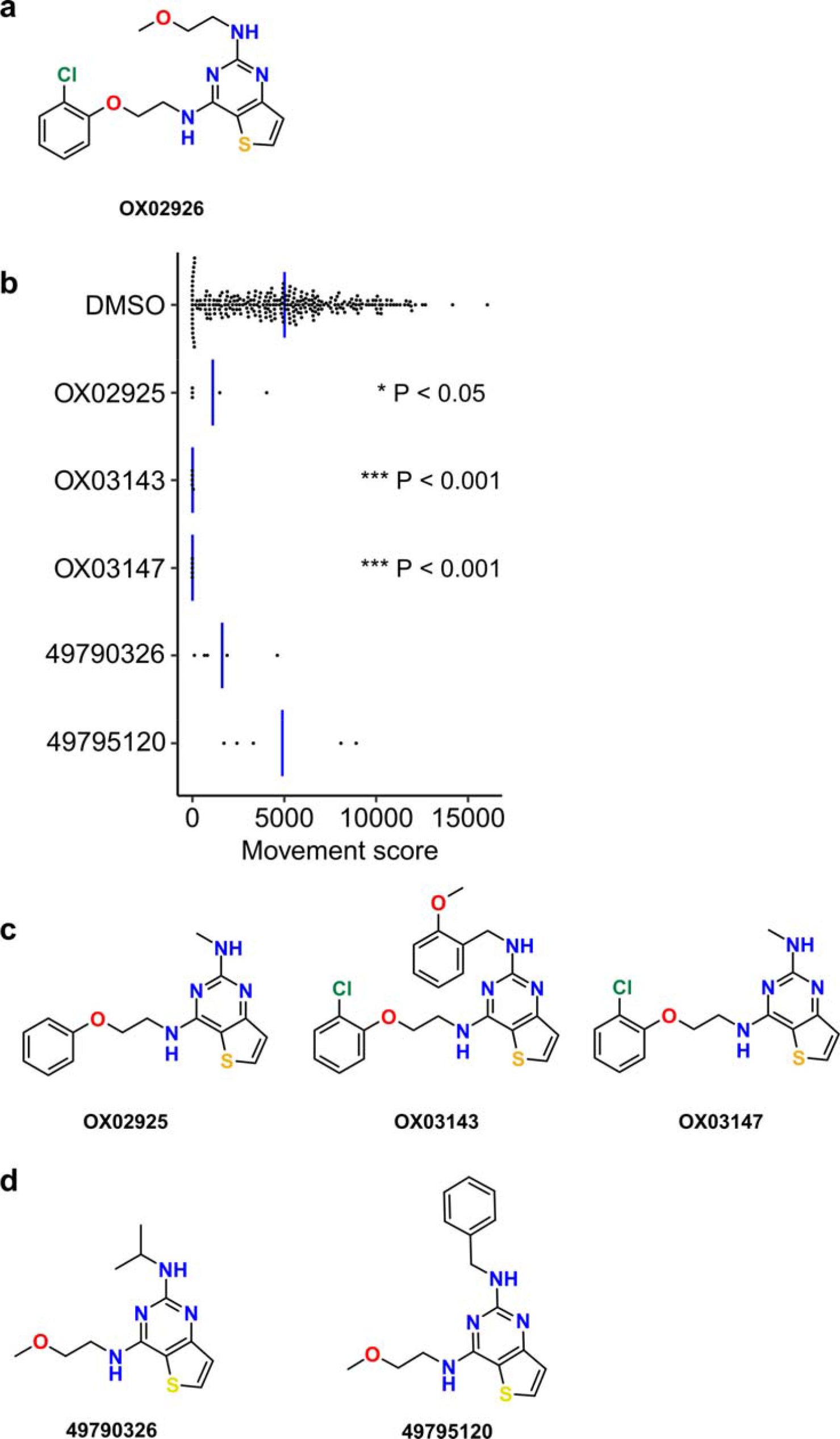
Identification of a diaminothienopyrimidine series from an *ex vivo T. muris* motility screen. **(a)** Structure of the hit compound, which was given the identifier OX02926. **(b)** Hit expansion by testing of structurally-related compounds using library material, assay concentration 100µM. Significance was determined by a two-sided Mann-Whitney test compared to DMSO-only controls, adjusted for multiple comparisons using the Bonferroni method (for test compounds n=5, each replicate on different assay plates, each point indicates one assay well). Blue bar indicates mean movement score. **(c)** Structures and identifiers of additional active compounds from this class. **(d)** Structures and PubChem CID accession numbers for the two compounds that were not significantly active in this assay.

### Resynthesis of active compounds

Having identified promising active DATPs from testing of DMSO solution samples of compounds, these were then resynthesised to obtain authentic, unambiguously characterised samples from which confirmatory screening could take place. Compound resynthesis is important since DMSO solution samples can degrade over time, and this often leads to so-called ‘false positive’ hits [19]. These compounds could be readily prepared in two steps from commercially available 2,4-dichlorothieno[3,2-*d*]pyrimidine **1**, via two sequential nucleophilic aromatic substitution reactions. Treatment of **1** with 2-(2-chlorophenoxy)ethylamine or 2-phenoxyethylamine gave exclusively monosubstitution affording **2a** and **2b** as a single regioisomer in 64% and 80% yield respectively. Subsequent displacement reaction at C4 gave authentic samples of **OX02925**, **OX02926**, **OX03143** and **OX03147** in 57 – 91% yield (Fig 2).

**Fig 2.**
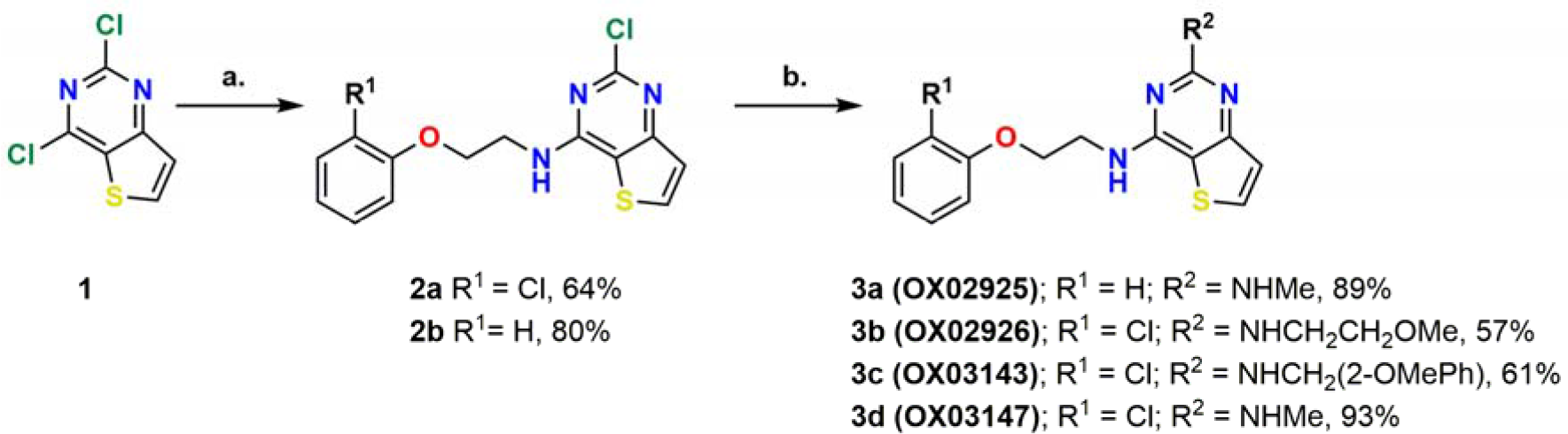
Synthetic route to putative hit compounds. **(a)** Substituted 2-phenoxyethan-1-amine (1.0 equiv.), DIPEA (2.0 equiv.), 1,4-dioxane, 80 °C, 3 hours. **(b)** Alkyl amine (10.0 equiv.), ^*i*^PrOH, 100 °C, 16-24 hours.

Activity of resynthesised diaminothienopyrimidines in the *T. muris ex vivo* adult motility assay The resynthesized hits were then tested in this screen and a concentration-response curve constructed, thereby confirming the anthelmintic activity of several examples of this structural class. (Fig 3, Table 1).

**Fig 3.**
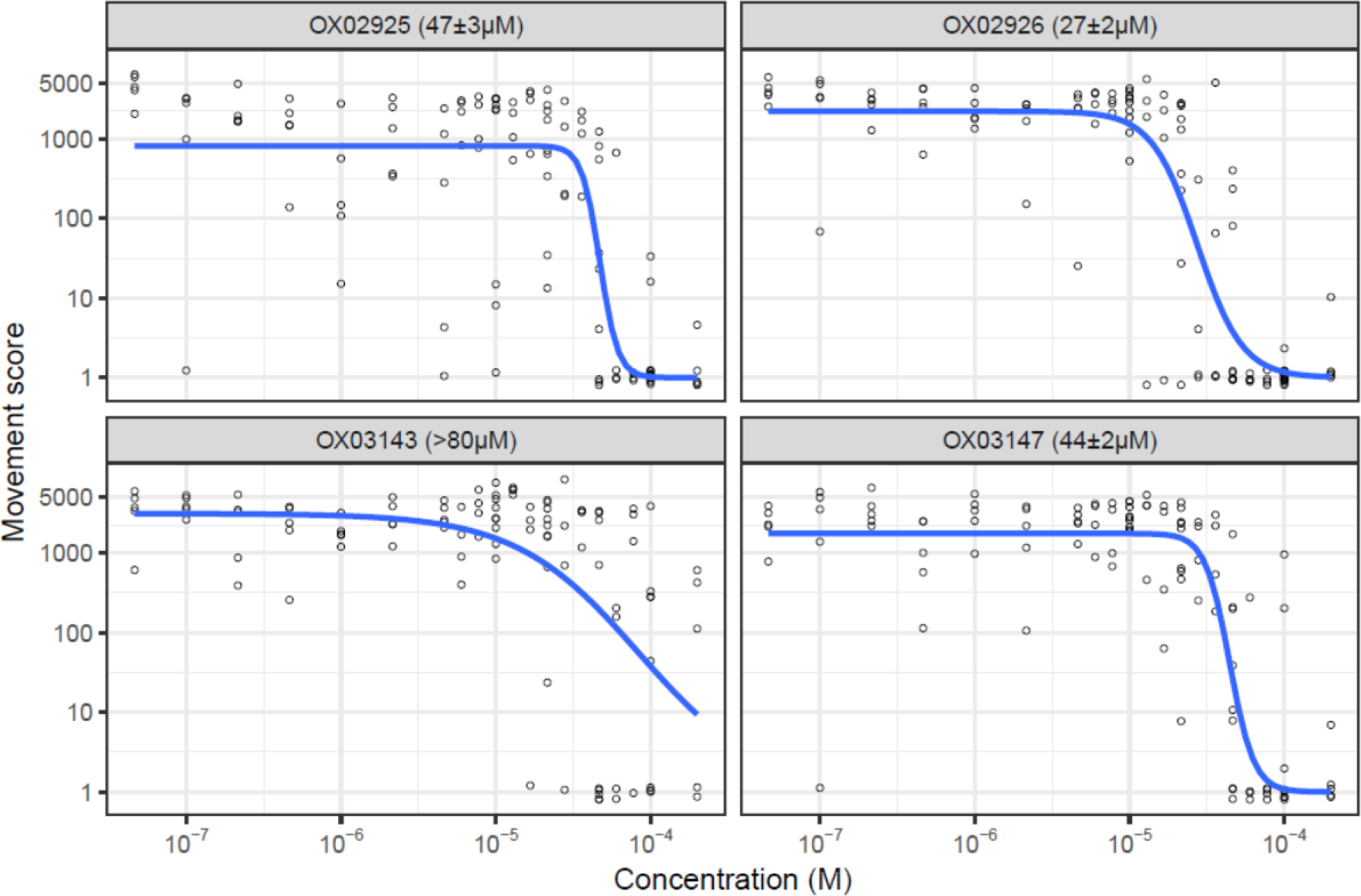
Concentration-response curves for resynthesized DATPs in the *T. muris ex vivo* adult motility assay. n=4 or 5 wells per concentration per compound, each replicate on a different 96-well plate using worms from different mice. Blue line indicates concentration-response curve fitted with the 3-factor log-logistic model using *drc* [16]. Figure in parenthesis indicates EC_50_ estimate ± standard error from this model. OX03143 did not clearly form a sigmoidal curve in the range of concentrations used in this assay so we report the EC50 estimate as > 80µM.

**Table 1.**
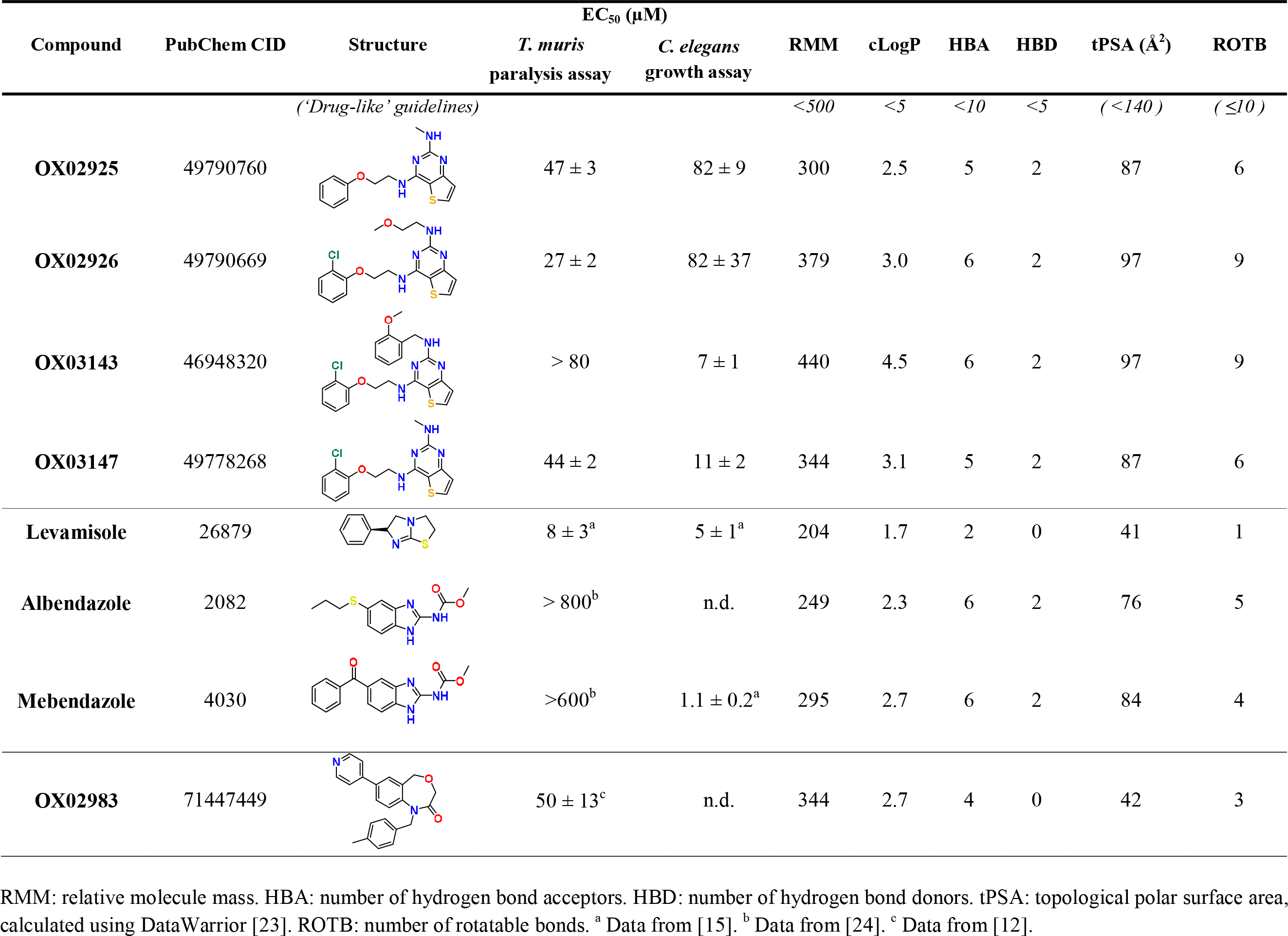
Properties and activities of resynthesized diaminothienopyrimidines, and other anthelmintics.

| Compound | PubChem CID | Structure | EC<sub>50</sub> (μM) |  | RMM | cLogP | HBA | HBD | tPSA (Å<sup>2</sup>) | ROTB |
| --- | --- | --- | --- | --- | --- | --- | --- | --- | --- | --- |
|  |  |  | *T. muris*<br>paralysis assay | *C. elegans*<br>growth assay |  |  |  |  |  |  |
| ('Drug-like' guidelines) |  |  |  |  | <500 | <5 | <10 | <5 | ( <140 ) | ( ≤10 ) |
| OX02925 | 49790760 |  | 47 ± 3 | 82 ± 9 | 300 | 2.5 | 5 | 2 | 87 | 6 |
| OX02926 | 49790669 |  | 27 ± 2 | 82 ± 37 | 379 | 3.0 | 6 | 2 | 97 | 9 |
| OX03143 | 46948320 |  | > 80 | 7 ± 1 | 440 | 4.5 | 6 | 2 | 97 | 9 |
| OX03147 | 49778268 |  | 44 ± 2 | 11 ± 2 | 344 | 3.1 | 5 | 2 | 87 | 6 |
| Levamisole | 26879 |  | 8 ± 3<sup>a</sup> | 5 ± 1<sup>a</sup> | 204 | 1.7 | 2 | 0 | 41 | 1 |
| Albendazole | 2082 |  | > 800<sup>b</sup> | n.d. | 249 | 2.3 | 6 | 2 | 76 | 5 |
| Mebendazole | 4030 |  | >600<sup>b</sup> | 1.1 ± 0.2<sup>a</sup> | 295 | 2.7 | 6 | 2 | 84 | 4 |
| OX02983 | 71447449 |  | 50 ± 13<sup>c</sup> | n.d. | 344 | 2.7 | 4 | 0 | 42 | 3 |

RMM: relative molecule mass. HBA: number of hydrogen bond acceptors. HBD: number of hydrogen bond donors. tPSA: topological polar surface area, calculated using DataWarrior [23]. ROTB: number of rotatable bonds. <sup>a</sup> Data from [15]. <sup>b</sup> Data from [24]. <sup>c</sup> Data from [12].

### Chemical properties of the hit series and synthetic suitability for further development

This class has ‘lead-like’ or ‘drug-like’ chemical properties [20], although it is important to note that in the contemporary medicinal chemistry literature this term is usually applied in the context of imparting oral bioavailability characteristics (Table 1). For agents targeting the gastrointestinal located *Trichuris*, minimal systemic exposure of the host is desirable and therefore it is critical to differentiate between the conventionally used terminology and parameters for ‘drug-like’ molecules, which affect solubility and permeability, compared to properties that would be relevant to agents targeting other body compartments. Recent literature has described this important caveat for non-peripheral CNS drugs [21], and indeed for anti-parasitic drug development [22]. Importantly, there is considerable scope for generating the large number of structural variants of the DATPs needed for the iterative improvement of compound properties during the downstream lead optimisation process.

### Active diaminothienopyrimidines block *C. elegans* development

Although we are focused on developing an anthelmintic with improved efficacy over existing drugs against *Trichuris*, activity across the nematode phylum is valuable, particularly as efficacy against economically significant agricultural animal parasites would make further development more economically viable.

We therefore wanted to test the activity of the DATP chemotype against the clade V nematode *Caenorhabditis elegans*. Using a quantitative development assay to measure the growth of synchronised L1 stage worms, we tested varying concentrations of the compounds to determine the concentration-response effects. As shown in Fig 4, all four DATP compounds were active in this assay with EC_50_ values from 7 – 87 µM.

**Fig 4.**
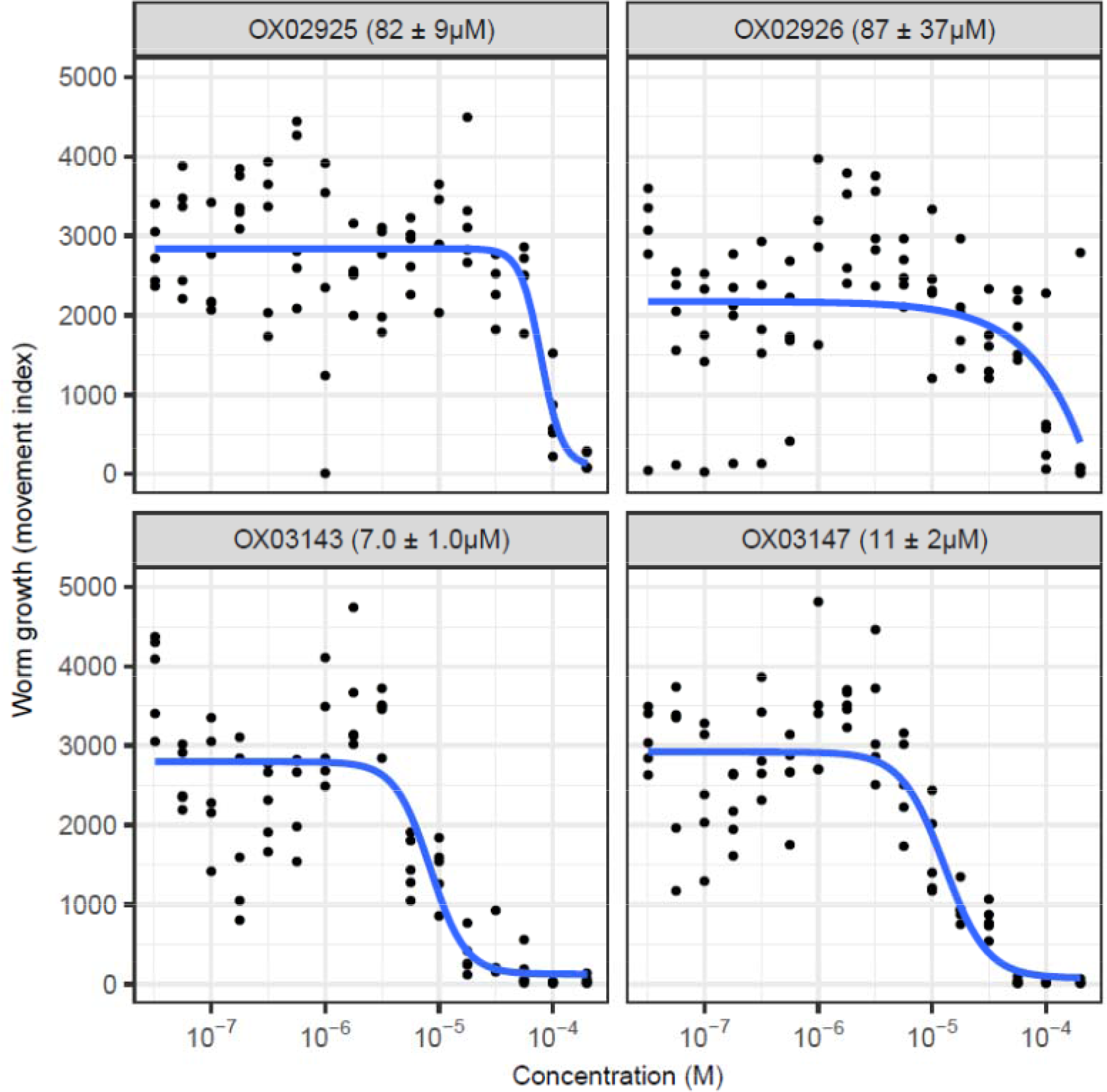
Concentration-response curves for resynthesized DATPs in the *C. elegans* growth assay. n=5 wells per concentration per compound, each replicate on a different 96-well plate. Blue line indicates concentration-response curve fitted with the 4-factor log-logistic model using *drc* [16]. Figure in parenthesis indicates EC_50_ estimate ± standard error from this model.

Interestingly, the DATPs display differing trends in activity between the *Trichuris* and *C. elegans* assays. At this stage we do not know whether this reflects different potency at the target or different patterns of drug access between the species, but the findings highlight the importance of screening against *Trichuris* in the search for novel anthelmintic agents targeting whipworm. The data from each of these assays as well as structural descriptors and Lipinski rule assessment for the four DATP compounds and other anthelmintics are summarised in Table 1. The leading member of the dihydrobenzoxazepinone class OX02983 is shown in Table 1 for comparison. EC_50_ values for the two series are currently in a similar range.

### Assessment of the cytotoxicity of the diaminothienopyrimidine series

It was critical to ensure that this series of compounds showed minimal cytotoxicity towards mammalian cells, and showed selective activity against the parasite. For example, gut cytotoxicity may result in the compounds having too narrow a therapeutic window. Selected examples of the DATPs were assessed for cytotoxicity using the mouse gut epithelial cell line CMT-93 (Table 2). Although, the DATPs exhibited increased *in vitro* cytotoxicity in these assays compared to the previously reported DHB series [12], an encouraging overall profile was exhibited for these early stage molecules. Furthermore, the nematode cuticle often limits drug access which reduces target engagement by small drug-like molecules [25,26]. This means that compound optimisation to improve uptake through the cuticle may be a fruitful route to improved anti-nematode selectivity, as well as improving the cytotoxicity profile.

**Table 2.** Summary of the cytotoxicity in a mouse epithelial cell line of the DATP series. Mouse CMT-93 rectal epithelial cells were used for this assay. Maximum tested concentration was 100 µM. *n*=8, error range (in parentheses) shows 95% confidence interval. EC_50_ values in the adult *Trichuris* paralysis assay are shown for comparison.

|  |  |  |  | 437 |
| --- | --- | --- | --- | --- |
| Compound | WST-8<br>$EC_{50}$ ($\mu M$) | Neutral red<br>$EC_{50}$ ($\mu M$) | Adult *Trichuris* paralysis assay<br>$EC_{50}$ ($\mu M$) | 438 |
| **OX02925** | 75 (48-124) | 29 (19-43) | 47 |  |
| **OX02926** | 43 (28-67) | 21 (15-31) | 27 | 439 |
| **OX03143** | 15 (9-26) | 5 (3-7) | >80 |  |
| **OX03147** | 37 (24-57) | 21 (14-30) | 44 | 440 |

It is interesting to note that the activity against *Trichuris* did not correlate with cytotoxicity, with the most cytotoxic compound (**OX03143**) showing the lowest activity in the *T. muris* adult paralysis assay, with an EC_50_ > 80µM. This suggests that either anti-*Trichuris* activity is distinct from cytotoxic action, or that differential drug access can be exploited to achieve differential host-parasite activity. Either possibility is encouraging and suggests that continued exploration and iterative improvement of the DATP structure might be anticipated to deliver a more potent anthelmintic with acceptable host toxicity.

### Activity of diaminothienopyrimidines against the infective egg stages of *T. muris*

Developing novel anthelmintics to disrupt the *T. trichiura* life cycle at the egg stage represents an exciting and complementary strategy to an oral therapy and is particularly attractive as *T. trichiura* eggs are highly resistant to extreme temperature changes and ultraviolet radiation, thereby remaining viable in the environment for many years [27]. We assessed whether the DATP derivatives were capable of affecting either infection establishment or embryonation of eggs. We first explored whether the compounds could alter the establishment of infection by soaking embryonated *T. muris* eggs in the test compounds for 14 days, washing the eggs and then determining infectivity both *in vitro* and *in vivo* (Fig 5a).

**Fig 5.**
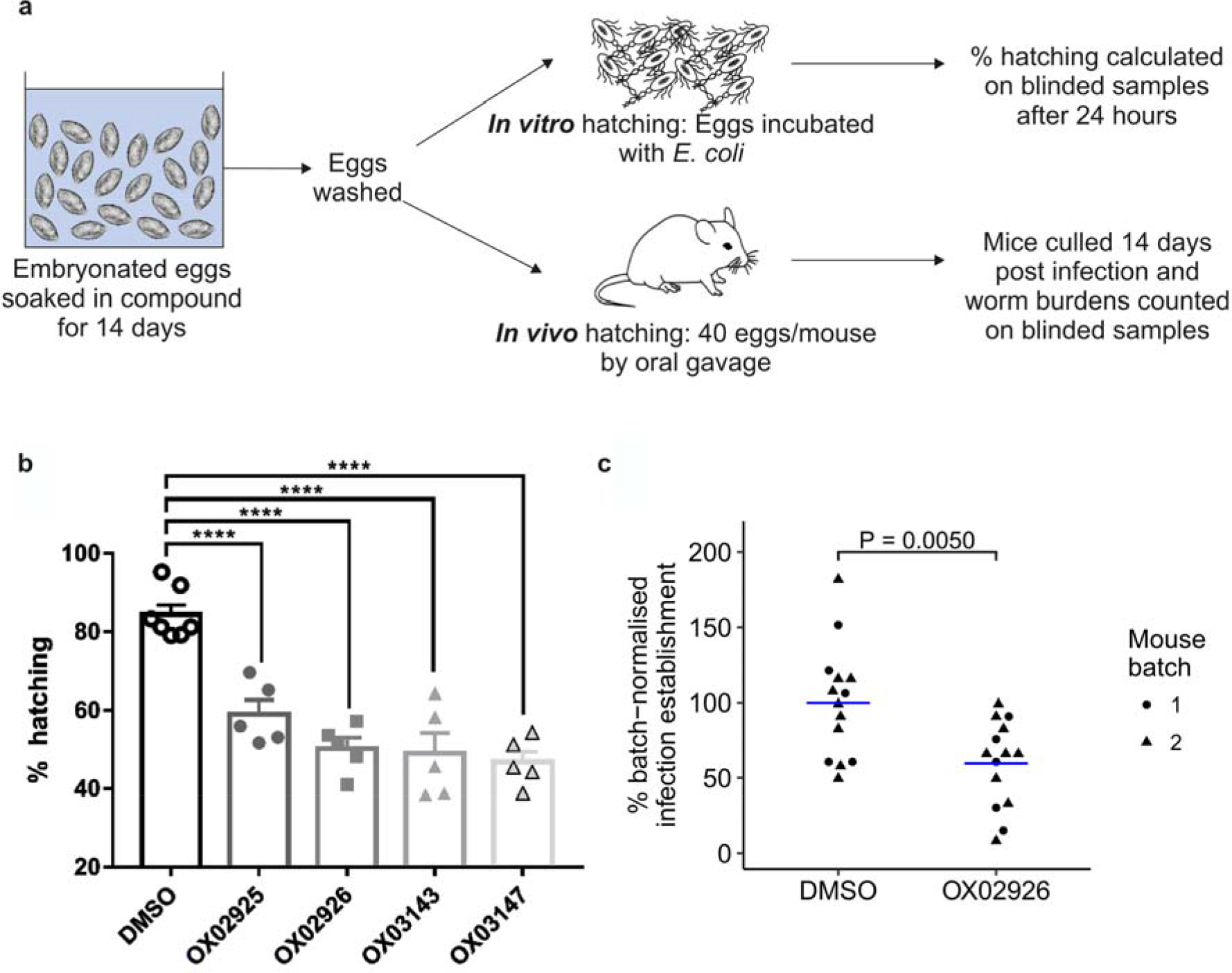
Reduced worm burden in mice given *T. muris* eggs that had been treated with diaminothienopyrimidines. **(a)** Embryonated eggs were soaked in compound for 14 days, washed in water and then used in either *in vitro* or *in vivo* hatching assays. **(b)** Treatment with DATPs reduced the ability of embryonated eggs to hatch in *E. coli* bacterial suspension after 24 hours. A one-way ANOVA showed a significant difference between treatment groups (F(5,26)=25.95 p<0.0001) with a post-hoc Dunnett’s compared to DMSO control (****= p<0.0001) *n*=7 (DMSO), *n*=5 (DATP compounds) **(c)** SCID mice were infected with 40 eggs and worm burden assessed at day 15 post infection. The experiment was carried out in two batches, with n=5 and n=9 mice respectively in each of the control and treatment groups. Data were normalised for each batch relative to the mean of the DMSO-only control group for that batch. Blue line indicates mean for each treatment group. A twoway ANOVA showed a significant effect of treatment [F(1,24) = 9.569, P = 0.00497] but no effect of batch [F(1,24) = 0.083, P = 0.77618] or interaction [F(1,24) = 0.083 0.77618]. A post-hoc Tukey HSD test showed that the **OX02926**-treated group was significantly different from the DMSO control group (P = 0.0050).

To determine effects on *in vitro* hatching, a protocol modified from that previously described [8] was established whereby eggs were induced to hatch when incubated in a culture of *Escherichia coli* at 37 °C. The results are summarised in Fig 5b. Strikingly all DATPs were capable of significantly reducing *in vitro* hatching compared to the DMSO control.

### Diaminothienopyrimidines reduce the ability of *T. muris* eggs to infect mice

To extend this finding, we selected **OX02926** to test in an *in vivo* hatching and infection establishment assay, as this compound showed both a significant decrease in *in vitro* hatching and a small standard deviation between samples. The eggs were soaked as for the *in vitro* experiment and SCID mice were infected with 40 treated eggs (**OX02926** or DMSO) by oral gavage. Egg infectivity was quantified at day 15 post infection by culling the mice and counting the number of established L2 larvae in the gut. All L2 larvae counted had a normal morphology as viewed under a dissecting microscope.

This experiment was carried out in two batches and the raw data are shown in the S2 Figure. Because variation in control worm establishment is commonplace in *Trichuris* infections due to natural variation in egg infectivity from a standardised egg number, we took the approach of normalising data for each batch relative to the mean of the DMSO-only control group for that batch. This allowed us to determine the effects of **OX02926** treatment (a full statistical description is given in the Methods section). The results are shown in Fig 5c. We used a two-way ANOVA (% batch normalised infection establishment ~ treatment * batch) to analyse the data. There was a significant effect of treatment [F(1,24) = 9.569, P = 0.00497] but no effect of batch [F(1,24) = 0.083, P = 0.77618] or interaction [F(1,24) = 0.083 0.77618]. We therefore conducted a post-hoc Tukey HSD test which showed that infection establishment in the **OX02926**-treated group was significantly different from the DMSO-treated control group (P = 0.0050). Treatment of eggs with **OX02926** was able to significantly reduce the burden of worms *in vivo* by an estimated 40%. This is likely to reflect reduced infectivity of DATP-treated eggs.

### Activity of diaminothienopyrimidines against the embryonation of *T. muris* eggs

The ability of the DATPs to alter the embryonation of *T. muris* eggs was investigated by soaking unembryonated *T. muris* eggs collected overnight from live adult *T. muris* in the test compounds at 26 °C for the duration of the embryonation process (56-60 days). During embryonation the first larval stage of the parasite develops within the egg shell (Fig 6a) from a ball of cells (Fig 6b). Treatment with the DATPs **OX02925** and **OX03147** resulted in a significant increase in the percentage of unembryonated eggs present compared to the DMSO control (Fig 6c). Importantly, although the other DATPs did not alter the percentage of eggs unable to undergo the embryonation process, the larvae that developed were atypical (Fig 6d-i). These atypical larvae were morphologically altered with the granules present within the larvae appearing less distinct.

**Fig 6.**
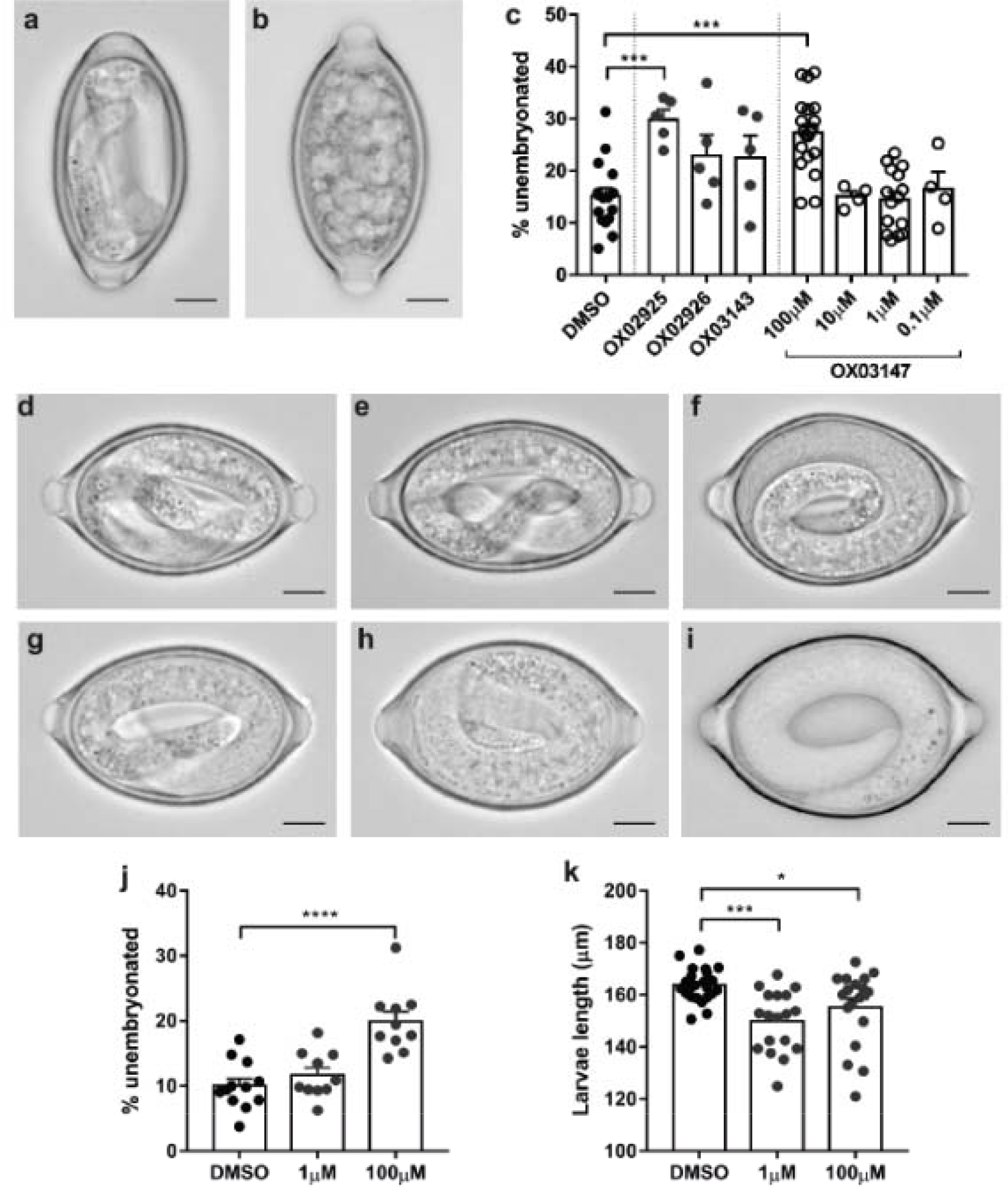
Unembryonated *T. muris* eggs treated with diaminothienopyrimidines have altered embryonation. Unembyronated eggs were soaked in 100 µM compound (unless specified otherwise) at 26°C (unless specified otherwise) for the duration of the embryonation process (56-60 days) and then embryonation determined and eggs imaged using an Olympus BX63 microscope. Scale bar indicates 10 µm. **(a)** Typical embryonated egg and **(b)** unembryonated egg. **(c)** treatment with DATPs increased the incidence of unembryonated eggs. Representative pictures of **(d)** DMSO, **(e) OX02925**, **(f) OX02926**, **(g) OX03143** and **(h) OX03147** 100 µM and **(i) OX03147** 1 µM soaked *T. muris* eggs. **(j)** Unembyronated eggs soaked in **OX03147** at room temperature for 56 days and embryonation determined. **(k)** Unembyronated eggs soaked in **OX03147** at 26°C for 56 days and larval length calculated using ImageJ.

As **OX03147** had the clearest phenotype with a significant increase in the number of unembryonated eggs, a concentration response study was performed to determine if an effect could be seen at lower treatment doses. Additionally, we repeated the experiment at room temperature to allow for more physiological conditions rather than the constant 26 °C utilised in the initial study to standardise conditions across experiments. Although the increased number of unembryonated eggs was only detected at the highest drug dose tested (100 µM) at both 26 °C and room temperature (Fig 6c, j) striking effects on egg morphology was detectable at concentrations as low as 1 µM with significant larval stunting observed (Fig 6k).

To determine if an effect on embryonation could be observed following a shortened drug exposure we soaked unembryonated eggs in 100 µM OX03147 at 26 °C for weeks 0-2, 0-3, 2-4 or 4-6 of embryonation. Although there was no increase in the proportion of unembryonated eggs observed in any treatment group (S3 Figure f) there were clear morphological alterations in the L1 larvae within the egg following exposure to OX03147 during weeks 0-3, 2-4 or 4-6 of the embryonation process (S3 Figure a-e). The most striking observation was the clear larval stunting observed following drug soaking from weeks 0-3 (S3 Figure c, g). A one-way ANOVA test showed a significant effect of treatment on larval length, F(4, 21) = 3.984, P=0.0147. A post-hoc Dunnett’s test showed a significant difference in the Weeks 0-3 treatment group compared to the DMSO-only control group (P = 0.0076). This appeared to phenocopy the effect OX03147 had when treated for the duration of the embryonation process at 1 µM (Fig 6k). Additionally, in the 2-4 week and 4-6 week groups, although larval length was not affected, there was evidence of structural alterations in the L1 larvae with a less distinct structure present and altered granulation within the larvae (S3 Figure d, e).

To the known range of applications of DATPs in medicinal chemistry we can now add anthelmintic activity. This study suggests they have significant potential for further development into dual-acting therapeutic agents for both the reduction of *Trichuris* egg infectivity, and embryonation in the environment. Thus, their actions on both the embryonated and unembryonated egg stages may enable a break in the parasite lifecycle.

## Discussion

Gastrointestinal nematode parasites remain a significant human health burden. Current anthelmintics lack efficacy and achieve low cure rates, threatening the targets set by the World Health Organisation for control of soil-transmitted helminths [2,28]. In particular, existing drugs have notably low efficacy against *T. trichiura*, the human whipworm. *T. trichiura* may be especially difficult to target as it inhabits the large intestine and is in part intracellular [29]. The metabolically active anterior of the worm, the stichosome, is buried in the host epithelial cells lining the gut, affording some protection from orally delivered anthelmintics.

### Diaminothienopyrimidines (DATPs), a new anthelmintic chemotype

We recently reported a small molecule screen for new anthelmintics targeting the gastro-intestinal (GI) nematode parasite *Trichuris muris* that identified the dihydrobenzoxazepinone (DHB) chemotype. The DHBs had not previously been ascribed anthelmintic activity [10]. Here, we describe a second class of novel anthelmintic, the diaminothienopyrimidines (DATPs). The potential for this early stage series is significant; their chemical synthesis is facile and lends itself to iterative optimisation, which will facilitate structural modifications aiming, for example, to increase local epithelial penetrance and hence improve efficacy during future development. Furthermore, their straightforward production imparts a favourable cost benefit aspect to the series.

### Other thienopyrimidines – their applications and targets

Thienopyrimidines have received much interest in medicinal chemistry as they are bioisosteres for purines, such as the nucleic acid components adenine and guanine. They are also related to quinazolines, an important class of kinase inhibitors, including gefitinib and erlotinib, which act by recognizing the ATP-binding site of the enzyme [30]. Thieno[2,3-*d*]pyrimidines are a particularly important scaffold, with many reported examples of protein kinase inhibitors, as well as inhibitors of dihydrofolate reductase, kainate receptor agonists, and α_1_-adrenoreceptor antagonists [31].

The thieno[3,2-*d*]pyrimidine scaffold found in the compounds reported in this study, has also been investigated. A series of 2-aryl 4-morpholino derivatives have been identified as phosphatidylinositol-3-kinase inhibitors [32], leading to the discovery of the PI3K inhibitor GDC-0941 (pictilisib) [33] and the dual PI3K/mTOR inhibitor DGC-0980 (apitolisib) [34]. The structures of these compounds are shown in the S4 Figure, in comparison with the 2,4-diaminothieno[3,2-*d*]pyrimidine OX02926. Pictilisib and apitolisib are under development as anti-cancer agents, have been tolerated in Phase I trials for solid tumors, and Phase II trials have commenced [35,36].

A series of 2,4-diaminothieno[3,2-*d*]pyrimidines have been described as orally active antimalarial agents [37], with activity in the low nanomolar range against *Plasmodium falciparum*. The structures of these compounds are shown in the S4 Figure in comparison with OX02926. This anti-malarial series was later improved by systematic modification giving improved antimalarial activity, but unfortunately continued hERG inhibition [38]. Whilst our DATP compounds have the same core scaffold as the anti-malarial series, they have different substituents, and in particular lack the 6-aryl substituent that is critical for anti-malarial activity and found in all compounds tested for hERG activity. However, the authors were able to demonstrate that hERG activity could be removed through modification of the C1 substitutuent, suggesting that this potential liability is not instrinsic to the 2,4-diaminothieno[3,2-*d*]pyrimidine core. We have not yet performed hERG assessment of our compounds, but this will form an important part of the future development of this series.

A series of 2,4-diaminothieno[3,2-*d*]pyrimidines has also recently been reported as active against the endosymbiotic bacterium *Wolbachia*, with potential use against filarial nematodes [39]. In neither the anti-malarial or anti-*Wolbachia* case is the molecular target of the compounds known.

### DATPs, their potential and route to a new anti-whipworm oral therapy

The major goal of our research is to develop a new oral therapy for trichuriasis, which could be widely used in mass drug administration programs leading to the eradication of human whipworm. Such an agent should have a substantially higher single-dose cure rate than the current drugs used in mass drug administration, albendazole and mebendazole. Drug development is long process, and recent work has defined a set of criteria, tailored to neglected infectious diseases, for progression in the hit to lead and lead optimisation stages [40,41]. Our DATP series members are early-stage compounds in the development process. The compounds meet almost all of the criteria for hit selection in neglected diseases, including confirmed activity with resynthesized material, dose-dependent *in vitro* activity, a tractable chemotype that passes drug-likeness filters such as the Lipinksi rule of five, and an established synthetic route of only two steps [40]. The most pressing weakness of the series is the small selectivity window for their activity against the parasite compared to cytotoxicity in a mammalian cell line. Improving this property for these early stage compounds must be a priority for future development. The DATP compounds also meet some of the milestones in the hit to lead process, particular in terms of drug-likeness and the exploitability of the structure, giving the ability to generate variants and establish the structure-activity relationship and hence improve potency and selectivity [41]. The *in vitro* activity of OX02926 in the adult whipworm motility assay (EC_50_ = 27µM, equivalent to 10.2µg/ml) also reaches the activity threshold for lead compounds that has been determined for drug development against the microfilarial nematode *Brugia malayi* [41]. In summary the DATP series are promising early-stage compounds with a number of lead-like features. Improvement of potency, together with an understanding of parasite/host selectivity and pharmacokinetic properties will be the focus of the next steps of development.

### Activity against the egg stage of *T. muris*

In addition to activity against the adult stage of whipworm, the DATPs were also able to significantly reduce egg hatching, both *in vitro* and *in vivo*. These data are in keeping with members of the DHB series, which also were able to inhibit parasite egg hatching. However, unlike the DHB series, we identified members of the DATPs that also significantly reduced the percentage of eggs embryonating *ex vivo*, with other members of the DATP series appearing to disrupt the embryonation process, resulting in defects in embryonic elongation and abnormal egg shape. *Trichuris* egg embryonation occurs gradually and the mechanism by which it occurs is currently a poorly understood process. A detailed characterisation of the morphological changes which occur with the *Trichuris suis* egg during embryonation has been described and other *Trichuris* species appear to undergo the same process. Once the unembryonated, unsegmented eggs are deposited, the two clear, nuclei-like areas move together and fuse. Cellular division then begins, initially occurring asymmetrically with two blastomeres of unequal size. The larger blastomere then divides again and then subsequently each blastomere divides in two until a blastula formed of many small blastomeres develops. The initial larval differentiation then occurs with the appearance of a motile cylindrical embryo, which gradually turns into an infective larva with its characteristic oral spear. The fully developed larva is no longer motile and is thought to be an L1 larva as no moult is observed within the egg [42]. The embryonation process is temperature sensitive. The effect of temperature on egg embryonation has been characterised in detail in recent years for *T. suis* eggs with the embryonation process accelerated at 30-32 ° C compared to 18 °C, with degeneration of the eggs rather than embryonation observed at higher temperatures (40 °C). At low temperatures (5-10 °C) no embryonation occurs, however once these eggs are then transferred to optimal embryonation temperatures normal embryonation proceeds [43]. Similar temperature sensitivity has been described for other *Trichuris* species including *Trichuris trichiura* with different species embryonating with different kinetics [44,45]. More research is required to understand the mechanisms behind this embryonation process, which may then allow an even more targeted approach to breaking the life cycle.

### Potential and feasibility of an environmental treatment

Humans become infected with *Trichuris* via a faecal oral route. Adult parasites in the intestine shed unembryonated eggs, which pass out with the faeces and embryonate in the external environment over a period of five weeks. Eggs can remain viable in the environment for many months [46]. Parasite eggs are only infective if fully embryonated upon ingestion. Thus, the ability of the DATPs to disrupt both the infectivity of embryonated eggs and the embryonation process itself suggests a potential environmental control to decrease *Trichuris* infection rates in the field without the need to develop and administer a new oral anthelmintic to the infected population.

In particular, it has been noted that the environmental pool of infectious eggs makes those individuals successfully treated, typically once or twice per year, in mass drug administration programs at risk of reinfection [47]. It has therefore been proposed that improvements in sanitation are required in addition to anthelmintic MDA. We suggest that an environmentally-acting, egg-targeting agent, potentially developed from our DATP series compounds, could play a complementary role to help break transmission in parallel with MDA and santitation improvements.

Clearly it is not possible to widely treat large areas of endemic regions with such an environmental control. Instead, we envisage the targeted use of DATPs in the environment at sites of high parasite egg density; these might include for example focusing treatment around pit latrines, as it is known that pit latrines may be a focal point of infection with a high concentration of eggs of soil-transmitted helminths [48]. In a study in Ethiopia, *Trichuris trichiura* prevalence was higher in communities with greater latrine usage (compared to field or yard defecation), suggesting that basic pit latrines may in some circumstances be ineffective at reducing infection [49]. However improved sanitation facilities generally, including pit latrines, ventilated improved pit latrines, and flush toilets, do reduce STH infection rates [47,50].

Such an egg-targeted agent should have a limited negative effect on the environment, have a suitable formulation for practical delivery, and be able to block egg viability at low concentration in the environment. The DATP series, which damage egg development and infectivity when applied at fairly high concentrations (1 to 100µM) for quite long periods of time (from 2 to 3 weeks to 60 days) show potential for developing such an agent. However these properties need to be improved during future development, while achieving an appropriate safety and environmental profile.

## Conclusions

In summary we report the discovery of a new class of anthelmintic, the DATPs, which possesses activity directed against adult stage *T. muris* parasites and the egg stage. Importantly, as a chemical series the DATPS are notable, since they are relatively facile to produce synthetically thereby presenting considerable scope for structural modifications to improve efficacy and deliver an optimised agent.

## Acknowledgements

Special thanks go to Roger Meadows for his help with the microscopy, and to Prof David Lomas and Dr Steven Buckingham for useful discussions during the course of this work.

## Supporting information captions

**S1 File. 1H NMR, 13C NMR and HRMS Spectra supporting the synthesis of the DATP compounds**

**S2 Figure.**
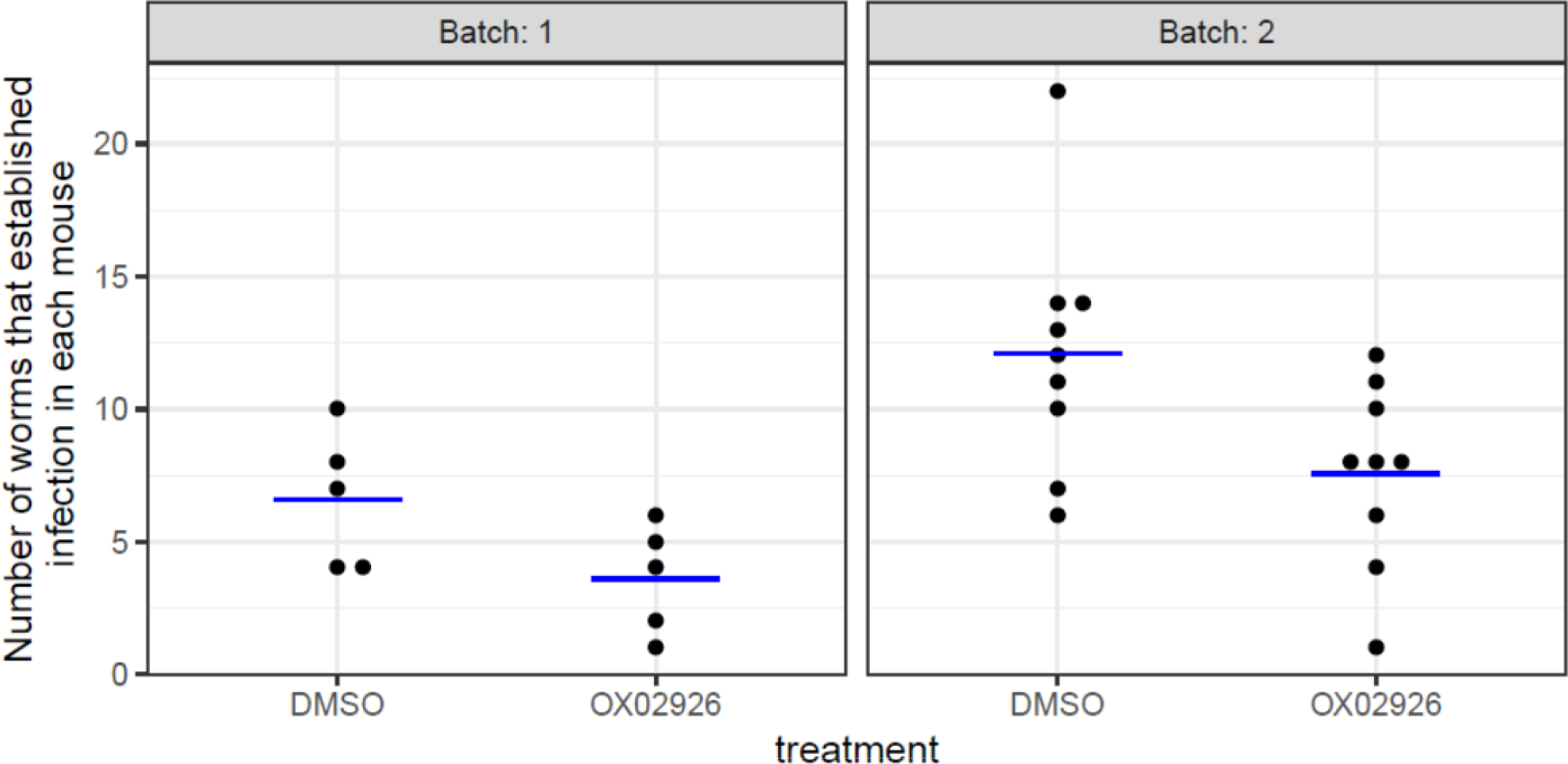
Raw data separated by batch for the in vivo hatching experiment. Each point indicates one mouse that has been infected with T. muris eggs that had been treated with deionised water plus 1% v/v DMSO (control) or deionised water plus 1% v/v DMSO and final concentration 100µM OX02926 for 14 days. Blue line indicates mean for each treatment group.

**S3 Figure.**
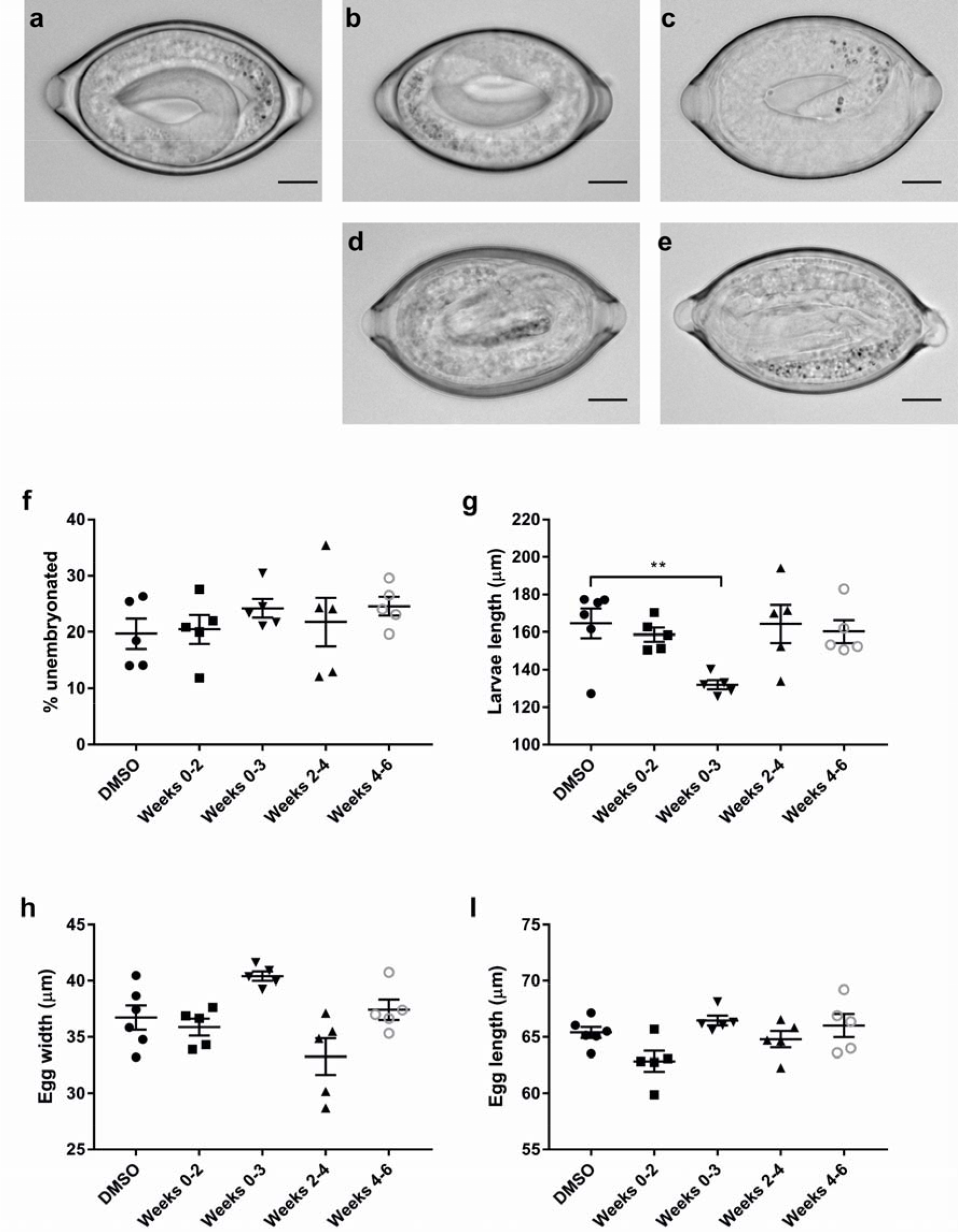
Unembryonated *T. muris* eggs treated with OX03147 for part of the embryonation process have altered morphology. Unembryonated eggs were soaked in 100 µM OX03147 at 26°C for the duration specified and then embryonation determined and eggs imaged using an Olympus BX63 microscope. Scale bar indicates 10 µm. Representative pictures of (a) DMSO, (b) OX03147 weeks 0-2, (c) OX03147 weeks 0-3, (d) OX03147 weeks 2-4, (e) OX03147 weeks 4-6. Following 56 days embryonation was determined (f) and larvae length (g), egg width (h) and egg length (i) calculated using ImageJ. ** Indicates P <0.01, one way ANOVA with post-hoc Dunnett’s test compared to DMSO control.

**S4 Figure.**
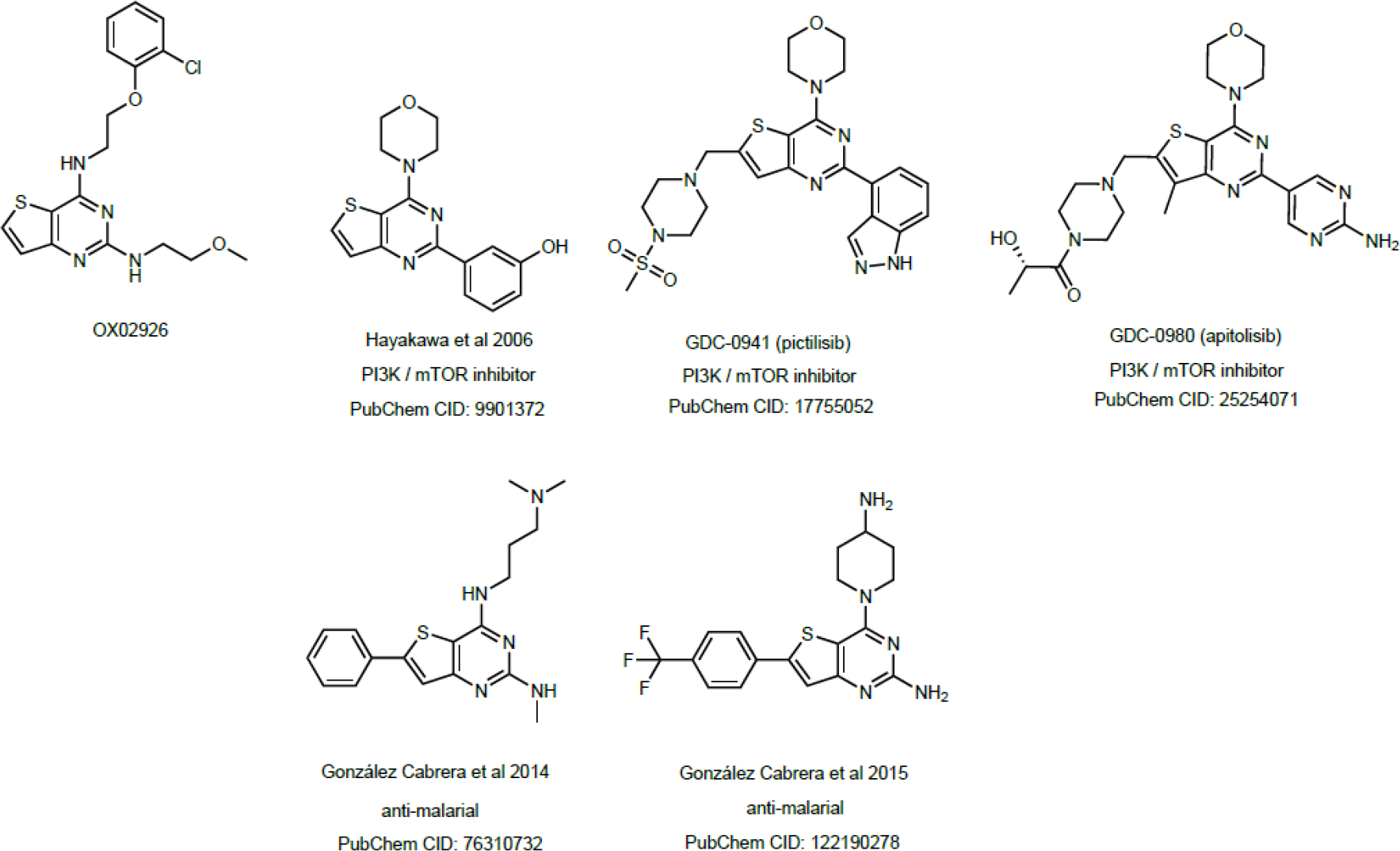
Structures of selected thieno[3,2-d]pyrimidine compounds in development compared to OX02926.

